# Simple degradable cyclodextrin polyester with chelator-based crosslinker for stent-based drug delivery

**DOI:** 10.1101/2021.04.29.442054

**Authors:** Kathleen Young, Audrey E. Lord, Grace E. Burkhart, Susan K. Kozawa, Nathan Mu, Horst A. von Recum

## Abstract

Cyclodextrins are a class of molecules which inclusion complexes with small hydrophobic drugs, and has historically been used to improve solubility and bioavailability of labile drugs in pharmaceutical applications. More recently, polymerized cyclodextrin has been applied in various applications as implantable drug delivery depots and as medical device coatings (e.g. polymeric hernia meshes) due to their ability to sustain and control drug delivery as well as prevent biofouling. Cyclodextrin polymers as coatings for metal medical devices, like screws or stents, is less explored; due to the high mechanical property mismatch between polymers and metals, a polymer coating is liable to delaminate easily, especially during device deformation. Novel methods for facilitating attachment to metal substrates have been explored, but coating longevity is still an issue, and these methods typically require the use of multiple reagents and complex methods. We report here the development and characterization of a cyclodextrin polymer with a chelator-based crosslinker with respect to appearance, chemistry, drug release profiles, erosion, pH-dependence. We found that increasing the crosslinking ratio (crosslinker:cyclodextrin) slowed down degradation and decreased drug loading as well. Drug release of the anti-restenotic drug sirolimus proceeded for over 4 weeks. The ability of the polymer to stably coat metal stents was verified, and the coating procedure is a simple, single step protocol.

## INTRODUCTION

Cyclodextrins (CDs) are a class of biomaterials widely used in drug delivery applications, from stabilizing labile drugs to enabling controlled drug delivery.^1^ CDs are cyclic oligosaccharides composed of 6-8 glucose units which form a ring-shaped molecule. The particular chemistry and shape of CD gives it a hydrophilic exterior dotted with hydroxyl groups as well as an interior hydrophobic pocket. CDs, with their hydrophobic cavities, are particularly adept at forming inclusion complexes with small hydrophobic molecules; in other words, CDs have affinity for many small hydrophobic drugs. CDs have historically been used in pharmacological applications to solubilize and improve bioavailability of drugs in vivo as well as to draw organic compounds out of solution, ^2,3^ and more recently as polymerized nanosponges, hydrogels, and particles for drug delivery.^4,5^ Polymerizing CD into highly crosslinked networks increase the likelihood of drug interactions with CD pockets, leading to increased drug loading and more linear drug release profiles compared to polymers without similar affinity.^4^

To take advantage of these controlled drug release abilities, CD polymers have been utilized as coatings on various types of medical devices, like orthopaedic screws^4^ and polymeric hernia meshes.^6^ Work on CD coatings for metal devices is uncommon, likely due to the high mechanical mismatch between CD polymers and metals, which can lead to the polymer easily delaminating during device deployment or day-to-day use. For deformable devices, like hernia meshes, which are made of polymers, these can be pre-treated with oxygen plasma, much like a cell culture dish, in order to facilitate covalent polymer attachment and significantly improve coating quality.^6^ The same cannot be said for metals; the high mechanical mismatch is a constant challenge for applying coatings, and metal surfaces usually require harsher treatments for surface activation or require an pre-treatment layer for the polymer coating to stably attach. The work that has been performed to obtain better attachment of CDs to stent surfaces has been focused on the addition of a pre-treatment polydopamine layer which attaches well to metals and provides a stable and degradable platform for crosslinking the CD polymers onto.^7^ A second method involves electrospinning a mix of CD polymers with chitosan to achieve a fibrous sheath-like coating that simply surrounds the abluminal stent surface.^8^ However, each of these methods involves the use of additional polymers and materials, which are potentially increasing the physical profile of the coating; minimizing stent profiles with thinner struts and thinner polymer coatings are generally desired.^9^ The added materials likely are also taking up valuable space which could instead be occupied by CDs which would increase drug loading efficiency as well as possibly extend drug release further by increasing drug residence time within the coating. We sought to develop a strategy where the CD polymer itself is able to stably attach to metal surfaces, without additional surface pre-treatments or coatings.

Ethylenediamenetetraacetic acid (EDTA)-crosslinked CD polymers have previously been synthesized from CD monomers, which forms highly crosslinked, insoluble CD nanosponges with a wide range of chemistries, which have been investigated for drug delivery as well as for pollutant (organics, metals) removal.^10,11^ Whereas these nanosponges are synthesized directly from the reaction of the crosslinking agent with CD monomers, our lab synthesizes CD polymers from CD prepolymers, which are small, soluble CD polymers that are lightly crosslinked with epichlorohydrin. By crosslinking the prepolymer, we obtain a highly crosslinked polymer that has an increased likelihood of inclusion complex formation with drugs, further slowing down and sustaining drug release.^12^ Previous methods of synthesizing these CD polymers involved the use of toxic crosslinkers,i.e. diisocyanates^4^ and diglycidyl ethers,^13^ and created non-degradable polymers. We hypothesized that crosslinking CD prepolymers with an EDTA-based crosslinker would retain the affinity while being able to chelate to metal stent surfaces for stable, near-covalent attachment. The EDTA-crosslinks produce ester bonds, so the resultant polymer should be degradable as well (**Scheme 1**).

In this paper we report the synthesis and characterization of a biodegradable CD polyester and is able to attach to metal surfaces due to its chelator-based crosslinker. We investigated several properties and functions of the new CD polymers, and varied the amount of EDTA to CD during synthesis to assess the effect of crosslinking ratio on polymer properties. To our knowledge, this paper is the first report of a stent/metal surface coating method for CD polymers which utilizes a simple, single step procedure and a single material for the coating.

## METHODS

### Materials

Soluble beta cyclodextrin polymer (prepolymer) which was lightly crosslinked with epichlorohydrin (CY-2009) was purchased from Cyclolabs (Budapest, Hungary). Ethylenediamenetetraacetic dianhydride (EDTA dianhydride), anhydrous dimethylsulfoxide (DMSO), and triethylamine were purchased from Fisher Scientific (Waltham, MA). Dextran (Mr 15000-25000) was purchased from Sigma (St. Louis, MO). Doxorubicin and sirolimus (Rapamycin) were purchased from LC Laboratories (Woburn, MA). MULTI-LINK VISION RX cobalt chromium (CoCr) Coronary Stent Systems (Abbott Vascular, Chicago, IL) were purchased from Esutures.com (Mokena, IL). 316L stainless steel foil shim was purchased from McMaster-Carr (Elmhurst, IL).

### Polymer Synthesis

CD polymers were synthesized using a modified procedure based on a previously published protocol.^10^ Briefly, EDTA dianhydride and beta-CD prepolymer were mixed in anhydrous DMSO with triethylamine at 25 degC. Crosslinking ratios of beta-CD to EDTA dianhydride of 1:6 and 1:10 were used to compare the effects of lower and higher levels of crosslinking on resultant polymer properties and performance. After reacting for 10 minutes, the polymer solution was vacuum filtered (0.2 um filter paper) while flushing with copious amounts of acetone to wash out excess unreacted substrate and solvent. Polymers were dried in the fume hood for 24 hours. For use in the following experiments, polymers were crushed with a mortar and pestle for 2 minutes into a fine powder. This synthesis procedure was also repeated for dextran, to make Dextran (Dex) polymers as a non-affinity control. Dextran was selected as the control due to its chemical similarity but structural difference to CD. Dextran and CD are both made of units of glucose, but dextran is a linear and branched polymer while CDs are cyclic molecules. Due to just this difference in structure, Dex does not possess the affinity for small hydrophobic molecules like CD and is therefore used as a non-affinity control.

### Scanning Electron Microscopy (SEM)

SEM was performed to characterize the appearance of EDTA-crosslinked polymers as well as polymer coatings. To prepare samples for SEM, double-sided carbon tape was first applied to SEM stubs. A small amount of polymer or a stent was gently pressed onto the tape. Stubs were tapped on the benchtop to remove excess polymer. Samples were sputter coated with 10 nm of palladium and imaged on a Helios Nanolab 650 SEM (FEI Company, Hillsboro, OR).

### Fourier Transform Infrared Spectroscopy (FTIR)

FTIR was performed to assess whether or not the synthesis reaction was successful; FTIR absorbance spectra would show characteristic peaks for functional groups and bonds present in analyte. To prepare samples for FTIR, each polymer tested was crushed into a powder with potassium bromide (in approximately 1:50 mass ratio) in a mortar and pestle. The powder was added to a die and pressed into a pelleted disk under the pressure of 8 tons in a hydraulic press for 10 minutes. The pellet was removed from the die. A Varian Excalibur series FTIR spectrometer (Bio-Rad, Hercules, CA) coupled with Varian Resolutions software was used to record IR absorbance spectra for all samples. First, 100 background scans with resolution of 4 were taken using an empty sample holder. Then sample pellets were mounted into the holder and read with 400 scans at a resolution of 4. Reactants (beta-CD prepolymer, dextran, EDTA dianhydride) and unreacted dry mixtures of reactants were also analyzed by FTIR for comparison.

### Drug Loading and Release Studies

To quantify drug release, polymers were first loaded with either a model drug, doxorubicin (DOX), or with sirolimus (SRL), and then subjected to longitudinal release studies in buffered solutions. DOX is a commonly used cancer drug that is red in color and fluorescent, allowing for both visual and quantitative evaluation of drug loading and release. SRL, also known as Rapamycin, is an anti-inflammatory and cytostatic agent that has commonly been used in drug-eluting stents. Polymers were incubated in high concentration drug solutions (2 mg/mL DOX in DMSO; 20 mg/mL SRL in DMF) on an end-over-end mixer for 3 days to load with drug. Then drug-loaded polymers underwent repeated washes with distilled water to remove excess drug and solvent. Drug-loaded polymers in DI water were frozen and then lyophilized for 2 days to obtain solid drug-loaded polymer powders. Drug-loaded polymers (20 mg DOX-loaded polymers; 10 mg SRL-loaded polymers) were added to 1 mL of release solution. For DOX release, 1X phosphate buffered saline was used; for SRL release, 0.1% Tween80 in 1X PBS was used to provide a hydrophobic sink.^13^ During release, samples in solution were stored on a shaking incubator at 37 C at 100 rpm. At specified timepoints, drug release vials were centrifuged, and then 700 uL of release solution were sampled without disturbing the polymer pellet. Seven hundred microliters of fresh, drug-free buffer were added back to the release vials to maintain infinite sink conditions. Samples were analyzed by ultraviolet (UV)/vis spectroscopy on a Synergy H1 Microplate Reader (BioTek, Winooski, VT). DOX release samples were analyzed by measuring fluorescence (excitation wavelength: 480 nm; emission wavelength: 600 nm) and comparing to a DOX standard curve to quantify DOX release. SRL release samples were transferred to a quartz-bottom microplate (Hellma, Plainview, NY) and were analyzed by measuring absorbance at wavelength 278 nm, then compared to an SRL standard curve to quantify SRL release.

### Erosion and Sensitivity to pH

To profile how quickly the CD and Dex polyesters degrade, polymer erosion over time was measured by weight. The behavior of the polymer in buffered solutions at a wide range of pH was simultaneously assessed to understand the polymers’ behaviors at different pH. For erosion in near physiologic conditions, PBS (pH 7.4) was used. An acidic Citrate-Phosphate (McIlvaine) buffer of pH 4.0 was made by mixing a solution comprise of 38.5% 0.2M disodium phosphate (28.38 g/L of disodium phosphate in DI water) and 61.5% 0.1M citric acid (19.21g/L citric acid in DI water).^14^ A basic carbonate-bicarbonate buffer at pH 10.4 was made by mixing a 0.2 % w/v sodium bicarbonate and 0.8% w/v of sodium carbonate in DI water.^15^ The solutions were autoclaved, and pH was remeasured to verify buffer stability. To set up the erosion study, 20 mg of each polymer were added to 1.5 mL Eppendorf tubes containing 1 mL of buffer solution, and 3 tubes were made up for each polymer in each buffer for selected timepoint (n=3), which ran up to 28 days. Eroding samples were stored on a shaking incubator at 37 C and 100 rpm. At each timepoint, 3 tubes of each polymer type in each buffer condition were removed from the incubator. Vials were centrifuged, the solution was removed, and vials were left open in a chemical fume hood until dry. Dried polymers were weighed, and the erosion rate was calculated simply as follows, using the nominal 20 mg as the initial weight:

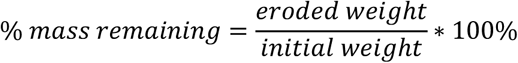

### Stent Coating and Surface Element Analysis

To determine whether the polymers could indeed coat metal surfaces, we performed a coating procedure, and then subjected coated stainless steel samples to surface element analysis. First the 316L stainless steel shim was cut into approximately 1 cm square pieces, and a single corner was crimped to provide a place to hold onto the thin samples without marking the rest of the surface. These were washed by sonicating in clean acetone and then clean MilliQ water, and then they were dried in a 100 C oven. Samples were cooled to room temperature and stored in petri dishes to prevent dust accumulation. To coat, we placed 10 mg of finely crushed polymers in 1 mL DI water in 1.5 mL Eppendorf tubes, and then up to 2 steel samples were added to each vial. The tubes were placed on an end-over-end mixer at room temperature for 24 hours. After the treatment, samples were washed copiously with MilliQ water (pointing bottle spout directly at surfaces) to remove any unattached polymers, and then they were air dried in a fume hood overnight. CoCr stents were cleaned and then coated with CD:EDTA 1:10 polymer in the same manner. A coated and an uncoated stent was then prepared for and imaged via SEM as described in the SEM methods section to determine coating appearance. Sample surfaces were analyzed via a Versaprobe 5000 X-ray photoelectron spectroscopy (XPS) Microprobe (PHI, Chanhassen, MN). Survey scans were obtained for each sample at 2 different spots near the center of the sample surface; for each of the 2 spots, 15 repeated scans were acquired. The different types of coatings analyzed were as follows: uncoated control, CD:EDTA 1:6 polymer, CD:EDTA 1:10 polymer (n=2). Survey scans returning the relative atomic % content of the most abundant surface elements in the energy range of 0-1100 electron Volts were obtained.

## RESULTS

**Scheme 1.**
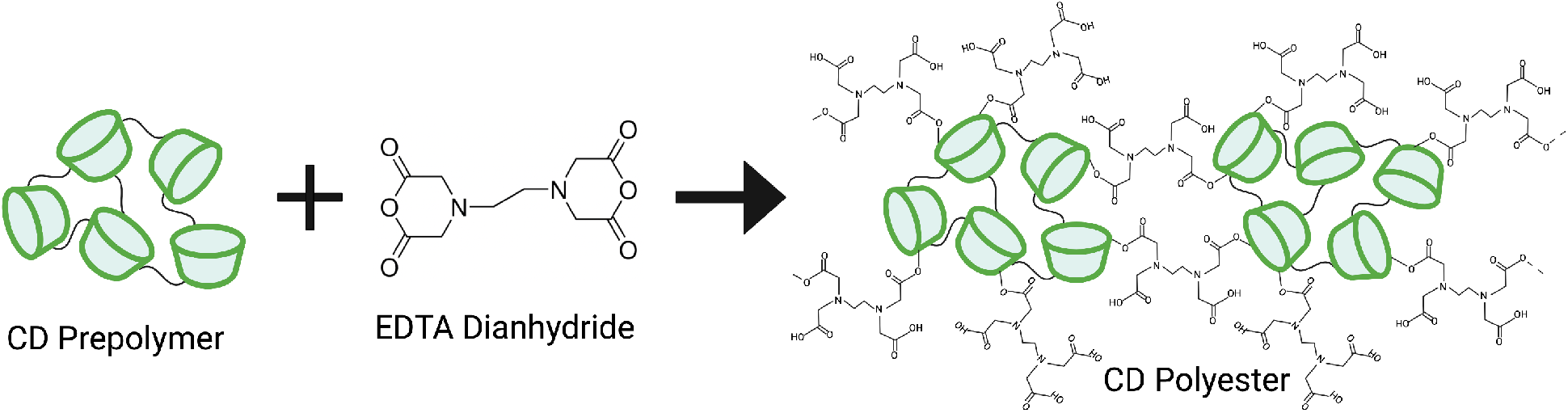
Particle synthesis and proposed CD polymer structure. Figure made with BioRender.com.

### SEM Images

SEM was performed to provide an initial look at the physical form of the newly synthesized polymers. From SEM images (**Fig. 1**), the crystallinity of the particles appears to depend on the crosslinking ratio – increasing the proportion of the EDTA-based crosslinker likely increases degree of crystalline structure. This is expected, CD prepolymers tend to be amorphous, and crosslinking with EDTA has previously produced polymers with crystalline structure.^10,11^ The Dex polymers appeared to be in relatively long, flat pieces (longer and larger than the CD polymers) despite the crushing procedure. This is likely due to the different starting polymer structures (CD prepolymer is amorphous, whereas Dex is linear and branched, and a little more salt-like). Unfortunately, particle size of the crushed polymers was not attainable from SEM images due to high levels of aggregation and clumping. To the naked eye, the polymers are white materials, and the crushed polymers are white powders.

**Figure 1.**
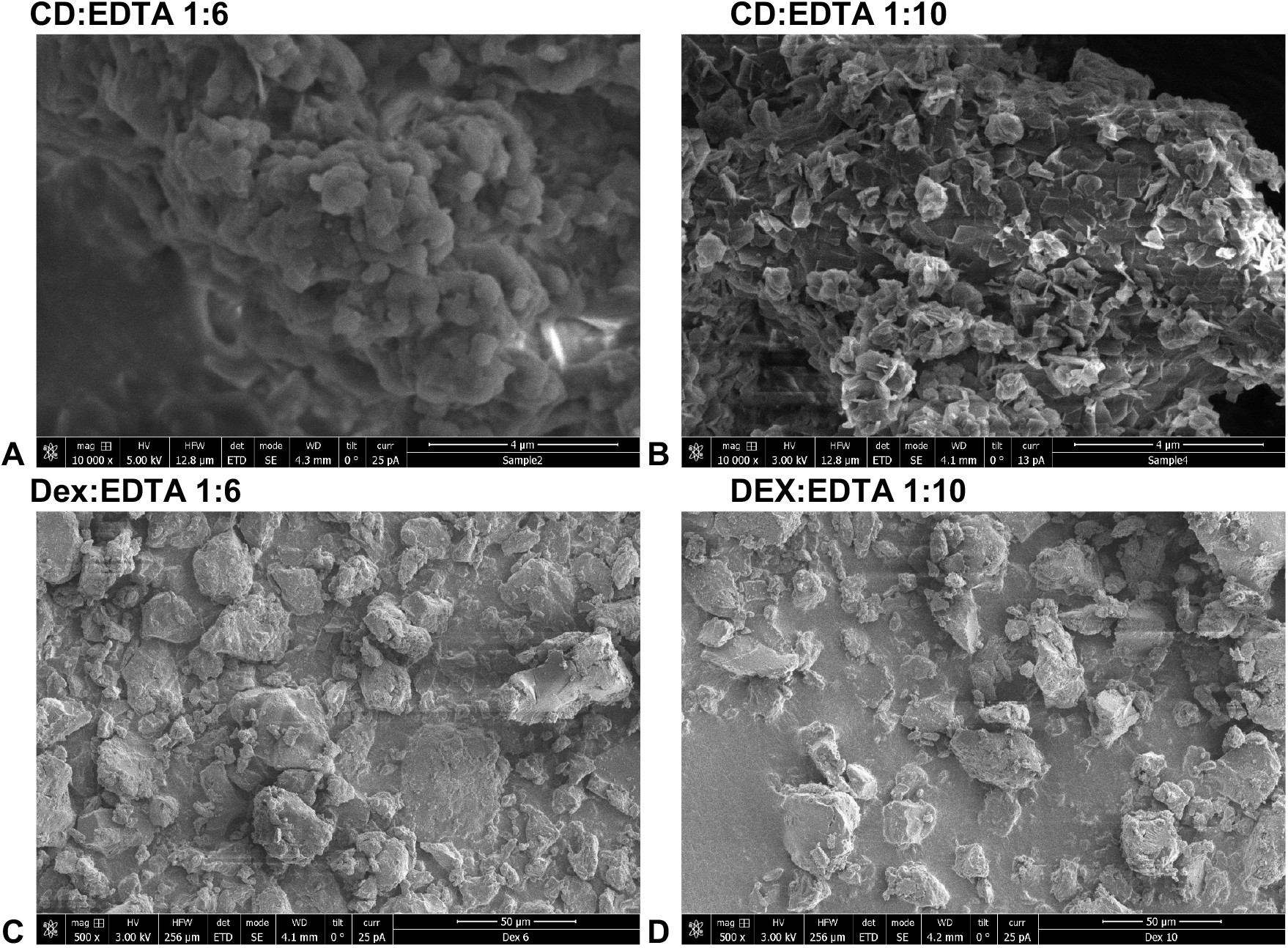
SEM images of synthesized polymer particles after crushing with mortar and pestle. **A**) SEM image of the less-crosslinked CD polymer (CD:EDTA ratio of 1:6) shows a relatively amorphous appearance of the polymer. **B**) SEM image of the more crosslinked CD polymer (CD:EDTA ratio of 1:10) shows a material with higher crystallinity than A, though with similar feature sizes (some particles less than 1 um in diameter). **C**) SEM of less-crosslinked Dex polymer (Dex:EDTA of 1:6) showing a relatively crystalline polymer (particles easily on the order of 20-30 um) which is much larger than the CD polymers In A and B. **D**) SEM of the more-crosslinked Dex polymer (Dex:EDTA of 1:10) shows increased crystallinity over the less-crosslinked version in C, with distinct and numerous flat facets on the polymer surfaces.

### FTIR Comparisons

To confirm that the synthesis reaction proceeded as expected, we used FTIR to identify newly formed, key functional groups in the polymers: ester bonds and carboxylic acids. Spectra for both types of CD polymers (1:6, 1:10) each showed a set of 3 peaks at around 1700, 1200, and 1000 cm^-1^ which are well-known to correspond to the C=O and the two C-O bonds present in an ester bond (**Fig. 2A**). Further, the spectra for CD polymers also show small peaks from 2800-2500 cm^-1^ corresponding to carboxylic acid (COOH) overtone and combination bands (**Fig. 2B**). Free COOH groups are formed due to the ring-opening of EDTA dianhydride during the reaction, as seen in the polymer schematic (**Scheme 1**). Similar sets of peaks showing the presence of new ester bonds and COOH groups were seen in the absorbance spectra for Dextran polymers as well (**Fig. 2C**,**D**), confirming that the synthesis reaction was successful. Absorbance spectra of unreacted dry mixtures (EDTA dianhydride + CD prepolymer; EDTA dianhydride + Dex) were also measured for comparison. These spectra lacked the sets of peaks and overtone bands for ester bonds and COOH groups (**Fig. 2E,F**), confirming that the mere presence of the reactants together does not produce the desired polymers.

**Figure 2.**
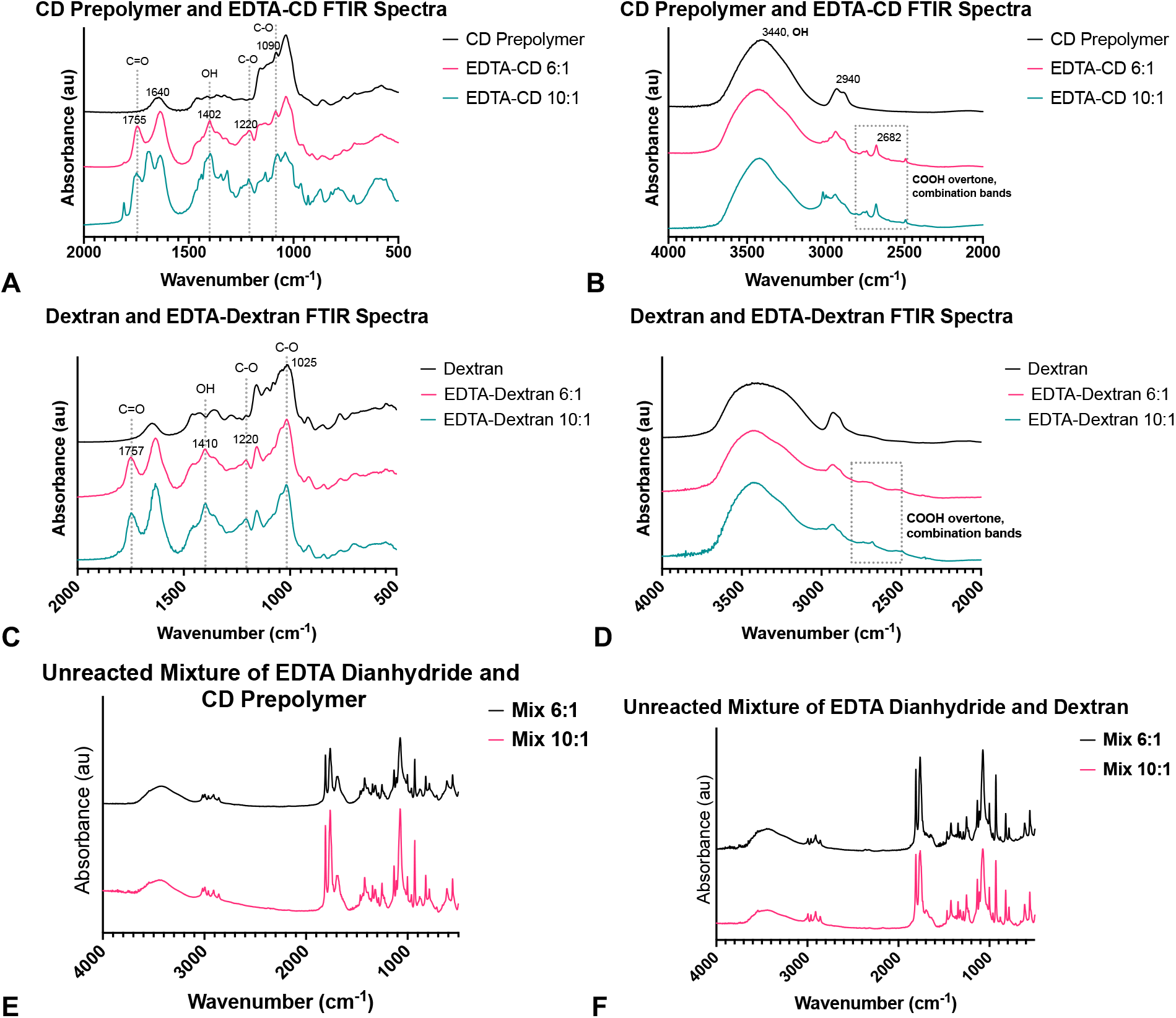
FTIR spectra of reactants compared to EDTA polymers. **A**) Comparison of CD polymers to the CD prepolymer precursor at lower wavenumbers. The EDTA-crosslinked polymers both showed characteristic peaks indicating ester bonds at approximately 1755, 1220, and 1090 cm^-1^.The OH bend highlighted at 1410 cm^-1^ is seen in CDs (polyols) as well as COOH groups. **B**) Comparison of CD polymers at a higher wavenumber range. The broad OH peak near 3500 is likely due to the alcohol groups of CDs. From approximately 2800-2500 cm^-1^, small peaks corresponding to COOH overtone and combination bands are present in only the EDTA-crosslinked polymers, not in the prepolymer. **C**) Comparison of Dex polymers at lower wavenumbers. They show the same characteristic peaks for ester bonds as the CD-EDTA polymers, as well as that same intensifying OH bend at 1410 cm^-1^. **D**) Dex polymers at higher wavenumbers, also showing the same COOh overtone and combination bands and a broad OH peak. **E**) The full absorbance spectra of unreacted mixtures of CD prepolymer and EDTA dianhydride lack the patterns seen in the reacted polymers. **F**) The full absorbance spectra of unreacted mixtures of Dex and EDTA dianhydride similarly do not show the characteristic peaks of ester bonds or COOH groups.

### Drug Release using DOX and SRL

We evaluated drug release from the CD and Dex polymers to understand the types of release profiles that can be achieved. We evaluated both levels of crosslinking on drug release, and we also tested two different drugs: a model drug, DOX, and the anti-restenotic drug, SRL. We performed DOX release studies in neutral PBS (pH 7.4), and SRL release studies were carried out in 0.1% Tween80 in PBS because SRL needs a more hydrophobic release sink to actually encourage release. **Fig. 3A** shows photos of DOX-loaded particles with CD:EDTA 1:6 on the left, and 1:10 on the right. The stark difference in the colors of these polymers shows a large difference in drug loading that is dependent on the crosslinking ratio during polymer synthesis. Less drug was loaded into the CD:EDTA 1:10 polymers, likely due to higher steric hindrance due to the presence of many more EDTA molecules within the polymers. release was sustained for up to 2 weeks (**Fig. 3B**). CD polymers also displayed sustained DOX delivery, not reaching more than 90% of drug release until day 8, compared to Dex polymers reaching 90% of the payload released by day 4. CD and Dex polymers sustained SRL release for more than 26 days (**Fig. 3C**). According to the % Drug Release scale, CD polymers appear to have a higher burst release and faster overall release compared to Dex. The maximum drug loadings of both DOX and SRL into all types of polymers assessed are compared in **Table 1**; this also shows that up to 10 times more drug can be loaded into CD polymers compared to Dex.

**Table 1.** Maximum drug loading, based on final cumulative release quantities.

| Polymer Type | Sirolimus (ug/mg polymer) | Doxorubicin (ug/mg polymer) |
| --- | --- | --- |
| Cyclodextrin, CD:EDTA 1:6 | 138.16 ± 5.41 | 34.52 ± 1.4 |
| Cyclodextrin, CD:EDTA 1:10 | 132.93 ± 3.12 | 24.32 ± 1.22 |
| Dextran, Dex:EDTA 1:6 | 138.71 ± 5.05 | 3.94 ± 0.24 |
| Dextran, Dex:EDTA 1:10 | 149.02 ± 13.45 | 3.40 ± 0.16 |

**Figure 3.**
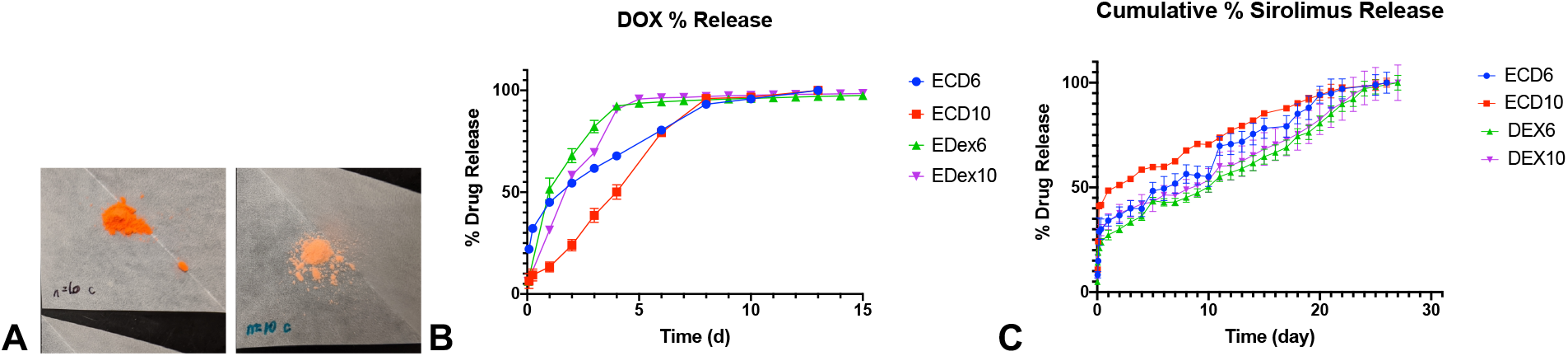
Drug release curves showing release of doxorubicin and sirolimus from polymers. **A**) Photos of DOX-loaded particles provide visual confirmation of difference in drug loading levels depending on crosslinking ratio. **B**) Cumulative percent release of doxorubicin shows total amount of doxorubicin released over time, with CD polymers sustaining DOX release over a longer period of time compared to the Dex controls. **C**) Cumulative percent release of sirolimus shows sustained release for at least 26 days. Error bars show standard deviation.

### Polymer Erosion

We performed erosion studies to quantify how the polymers degrade/erode over time. As these are polyesters, environmental pH was likely to impact degradation rates. We performed degradation in acidic, neutral, and basic buffers (pH 4.0, 7.4, 10.4 respectively). Release in acidic and neutral buffers was similar for all types of polymers tested over a period of 3 weeks (**Fig. 5 A-D**). At high pH, Dex polymers degraded almost immediately, whereas CD polymers did not degrade until 10-15 days, showing the better stability of the EDTA-crosslinked CD polymers.

**Figure 4.**
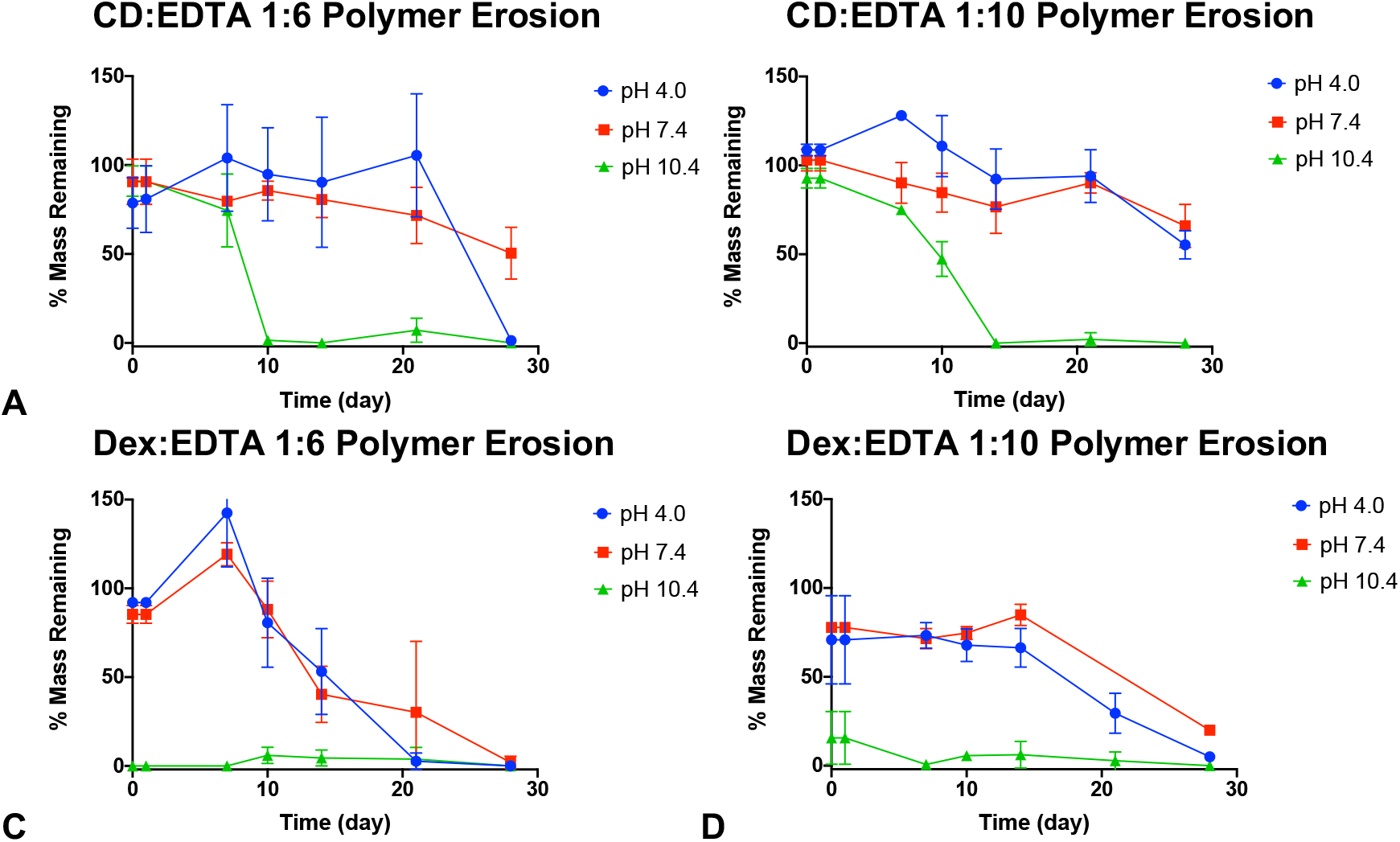
pH dependence of erosion of CD and Dex polymers measured by % mass remaining over time. **A**) Erosion of CD:EDTA 1:6 polymers, **B**) CD:EDTA 1:10 polymers, **C**) Dex:EDTA 1:6 polymers, and **D**) Dex:EDTA 1:10 polymers.

**Figure 5.**
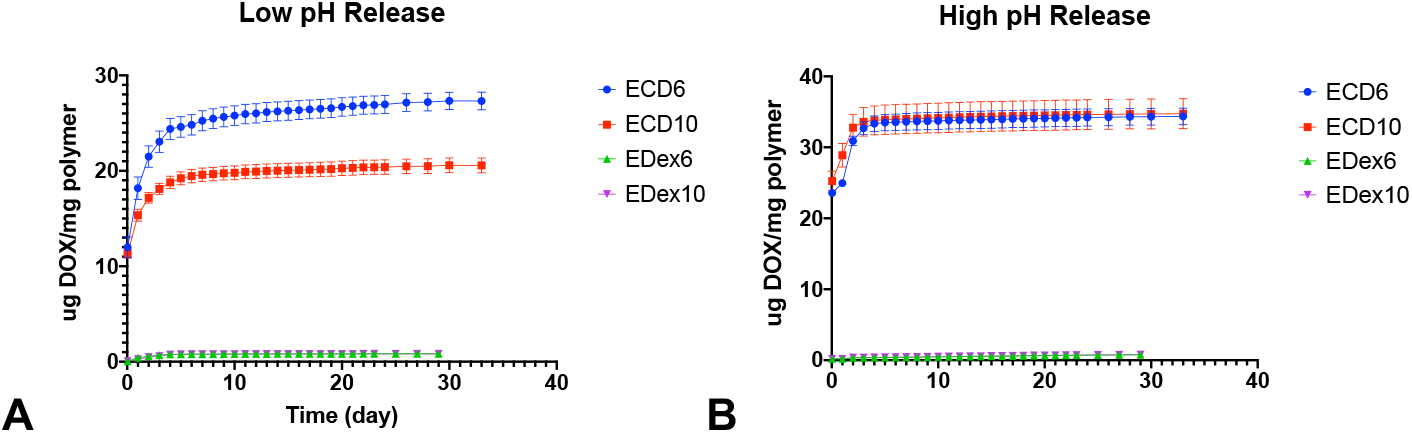
Drug release in acidic (pH 4.0) and basic (pH 10.4) buffers. **A**) DOX release from CD and Dex polymers in acidic/low pH buffer. Here CD shows release profiles which are similar to that in neutral PBS buffer in Fig. 3B. **B**) DOX release from CD and Dex polymers in basic/high pH buffer shows high burst release from all polymers, which reach 100% of drug released within 5 days.

### Drug Release in Acidic and Basic Buffers

To evaluate the effect of environmental pH and varying degradation rates on drug release, we performed drug release studies in low and high pH buffer solutions (citrate-phosphate buffer at pH 4.0 and carbonate-bicarbonate buffer at pH 10.4, respectively), using the model drug DOX. Daily drug release aliquots were taken for 23 days, then sampling frequency was reduced to once every 2-3 days. Drug release is reported as the mass of released DOX divided by the mass of the polymer at the start of each study to account for different initial polymer masses. At low pH, CD polymers showed a higher amount of drug release as well as more sustained release compared to Dex polymers (**Fig. 5 A**), likely owing to the higher drug loading capacity in CD. In high pH, all polymers appeared to catastrophically degrade, leading to high and immediate burst release in both CD and Dex (**Fig. 5B**), resulting in the majority of active release occurring within the first 5 days.

### Stent Coating Surface Analysis

Finally, we evaluated the ability of the CD polymers to form coatings on 316L stainless steel surfaces as well as on CoCr stents (Abbott). We imaged an uncoated and a CD polymer-coated CoCr stent. The uncoated stent surface is smooth and largely free from defects (**Fig. 6A**). In contrast the coated stent clearly has a bumpy texture with features on the order of 20 um in diameter, showing the presence of polymer on the stent surface despite vigorous washing with water (**Fig. 6B**). XPS was performed to acquire surface element composition of untreated and coated 316L stainless steel samples. XPS showed the expected increase in carbon (C1s) and nitrogen (N1s) content; the sources of these changes are the carbon in the polymers and the nitrogen in EDTA (**Fig. 6C**). There was a decrease in oxygen content, which likely is due to the presence of a passivating metal oxide layer at the surface of stainless steels. As the coating is applied and the steel surface is covered up, the relative amount of oxygen decreases. The relative amounts of iron (Fe2p3) and chromium (Cr2p3) captured by the XPS decreased as expected as well, for similar reasons to oxygen. Overall, CD:EDTA 1:10 polymer coating produced the greatest change, although this change was not significant (n=2). As stated above, we hypothesized that EDTA-crosslinked CD polymers are capable of chelating to metal surfaces due to the presence of EDTA crosslinks (which can still chelate metals out of solution^11^) and linked EDTA molecules which are not fully crosslinked between CD molecules (the 3 EDTA “arms” pictured chelating the steel surface in the schematic in **Fig. 6D**).

**Figure 6.**
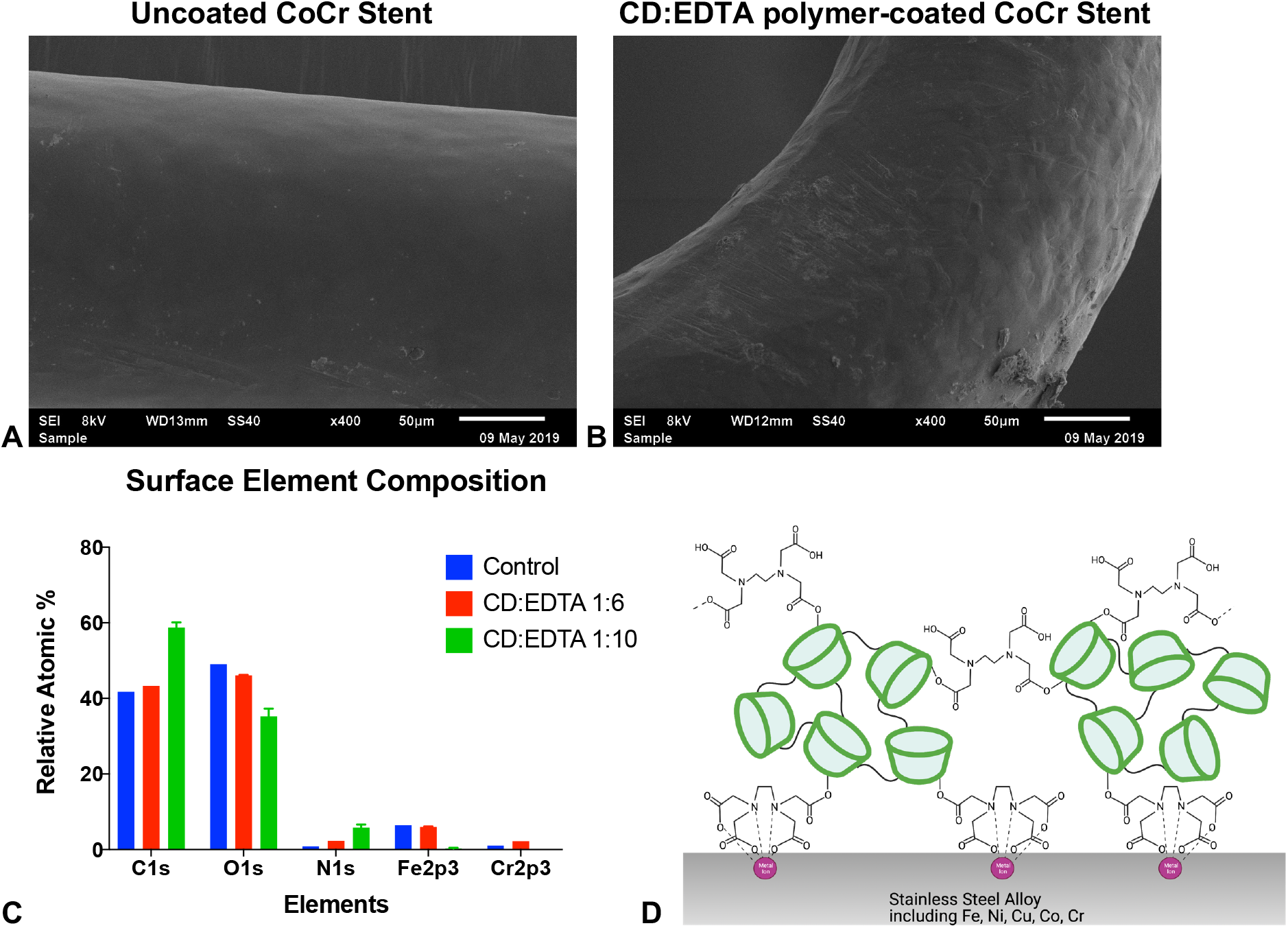
Preliminary evaluation of CD polymers as stent coatings. **A**) SEM image of uncoated, untreated CoCr stent. **B**) SEM image of CD:EDTA 1:10 polymer-coated CoCr stent; the texture shows the coating presence. **C**) XPS results showing surface element composition of 316L stainless steel samples (n=2). **D**) Schematic of how we theorize the EDTA-crosslinked CD polymers attach stably to metal surfaces via chelation. Figure made with BioRender.com.

## DISCUSSION

To summarize the main results presented in this paper, we characterized the chemical and physical make-up of this new type of biodegradable affinity CD polymer, and demonstrated proof-of-concept coating on metal and metal stent surfaces. FTIR confirmed the success of polymer synthesis. The insolubility of the polymers in aqueous buffers also signified successful polymer synthesis. We performed drug release studies with DOX and SRL, finding DOX release for 2 weeks and sustained SRL release for over 4 weeks. Erosion studies showed that CD polymers likely take much longer than 4 weeks to fully erode in neutral buffered solution. Polymer erosion was hugely accelerated in high pH buffer; comparatively, low pH buffer appeared to preserve polymer longevity. This pH-dependent erosion behavior follows the trends which are typical of degrading polyesters.^16^ This also points to the potential use of this polymer in low pH conditions.

We selected two crosslinking ratios of beta-CD to EDTA dianhydride (1:6, 1:10) to assess across all experiments in order to determine the effect of crosslinking ratio on polymer properties. Increasing the crosslinking degree slows down the erosion and degradation rate and also decreases the overall amount of drug loading, likely due to steric hindrance. Interestingly, our data shows that increasing the crosslinking ratio beyond 1:6 results in concomitantly increased crosslinking degree (more intense peaks in FTIR). Previous work on EDTA-crosslinked beta-CD nanosponges actually showed that the maximum degree of crosslinking occurred at a ratio of 1:6 CD:EDTA, likely due to steric hindrance during synthesis.^17^ Because we applied the polycondensation reaction to EDTA dianhydride with CD prepolymers, we did not see a similar effect limiting crosslinking degree. CD prepolymers are much larger than monomers (tens of kDa compared to approximately 1 kDa), so the crosslinking/reactive capacity of the CD prepolymers likely was not saturated out with the ratios used. In future work, properties of the polymers synthesized from CD prepolymers as precursors should be assessed. Based on the observation that the more crosslinked CD polymer eroded more slowly, degradation likely can be significantly slowed by increasing the crosslinking ratio. The non-affinity control Dex polymers created in this study would also benefit from substantially increasing the crosslinking ratio, because the Dex precursors (15-25 kDa) likely are also so large that the resultant polymers likely are not densely crosslinked.

A preliminary stent coating study was performed which confirmed the ability of these EDTA- crosslinked polymers to remain attached to metal and metal stent surfaces. In **Fig. 6D**, we present a hypothetical schematic showing how the polymer likely utilizes chelation as a mechanism to attach to metal surfaces. In this figure, the EDTA linkers shown interacting with the metal surface are not fully crosslinked between two CD molecules; we expect that it is more energetically favorable for an EDTA molecule with 3 free arms to fully form coordinate bonds compared to a crosslinked and restrained EDTA which only has 2 free arms. We also expect that the majority of EDTA crosslinked will be formed in the interior of the CD:EDTA polymers and will simply not have physical access external metal surfaces.

XPS data likely indicates that the CD:EDTA 1:6 polymer coating is thinner than the maximum penetration depth of the XPS probe, since the relative amounts of iron and chromium (metal substrate) are similar to the control. For the particular XPS instrument model used, the max probe depth is between 5-10 nm below the sample surface. Because the metal elements are essentially absent in the CD:EDTA 1:10 polymer coating samples, this indicates that these coatings are at least as thick as if not thicker than the probe depth of 5-10 nm, giving us an estimated coating thickness of 5-10 nm. However, a low sample size was used for XPS, which does not allow us to draw any statistically significant findings from this data.

Interestingly, XPS did show that the control, uncoated 316L stainless steel sample surface was comprised of more than 40% carbon, whereas a low corrosion stainless steel like this should have less than 1 or 2% carbon content. This is likely due to organic contamination which was not sufficiently removed during the washing steps. The washing procedure was consistent across all samples though, i.e. each steel sample had the same organic contamination, so the changes in carbon content observed are accurate.

Referring back to the introduction, we claimed to create a new coating method which is much simpler compared to the existing strategies used to attach CD polymers to stent surfaces. Here we report a coating method which is essentially a single incubation step with the polymer at room temperature in water, followed by a wash which is also in water. This simple method can be easily applied to metal medical devices of different shapes and sizes, not just coronary stents. The coverage and thickness of the coating can be tuned simply through adjusting polymer synthesis parameters (type of CD prepolymer, CD to EDTA crosslinking ratio).

We acknowledge several limitations of this study. First, for the in vitro work, we used common pH buffered solutions, which do not recapitulate the complex biological environment. The one exception to this is the SRL drug release, where 0.1% Tween80 in PBS was used as a more physiologic release medium. However, it is possible that even the addition of Tween80 was not sufficient to provide a sufficient hydrophobic sink to match infinite sink conditions. To address the zero-order release profiles of SRL from both CD and Dex polymers, it is likely that the SRL quickly reached the limit of solubility in the release medium due to the solution not being hydrophobic enough and that this is the cause of the near identical release from both the affinity and non-affinity polymers. Beta-CD was selected for use in this study due to its greater compatibility with SRL. However, DOX has more affinity with gamma-CD compared to beta-CD^18^, so synthesizing EDTA-crosslinked polymers from gamma-CD prepolymers should hypothetically show greater DOX loading and more sustained release due to higher affinity. The low sample size of XPS data does not allow us to draw

Future work in developing this polymer as a drug-eluting stent coating will have to involve the use of other varieties of CD and drug combinations. The optimal combination of CD size and drug should be selected to maximize the effects of affinity-based drug delivery. As previously mentioned, CD polymers with much higher crosslinking degrees should be synthesized and evaluated for drug delivery and degradation properties to understand how far this polymer can be tuned. Further assessment of the use of the CD polymers as drug-eluting stent coatings are needed; a drug delivery study from EDTA-crosslinked CD polymer coated stents is currently under way with more than 40 days of progress and continuous drug release (data not shown). Testing the coating durability, longevity, hemocompatibility, and cytocompatibility are all necessary components of drug-eluting stent coating development as well.

## CONCLUSION

In summary, we developed a new version of EDTA-crosslinked affinity cyclodextrin polymer and demonstrated its use in drug delivery as well as its ability to sustain delivery compared to non-affinity controls. We obtained drug delivery of doxorubicin for about 2 weeks and sirolimus for over 4 weeks, and we also observed erosion profiles characteristic to polyesters. Increasing crosslinking decreases drug loading to a degree and also slows down degradation, likely due to the addition of more crosslinks which take time to degrade. This work also offers a simple alternative method for creating a well-attached drug-eluting cyclodextrin polymer coating on metal stent surfaces, positioning this polymer as a promising drug-eluting stent coating material.

## ACKNOWLEDGEMENTS

This material is based upon work supported by the National Science Foundation (NSF) Graduate Research Fellowship under Grant No. 1937968 (KY) and the NSF Summer Research Experience for Undergraduates (GEB). The authors would also like to acknowledge the use of facilities at the Swagelok Center for Surface Analysis of Materials at CWRU, and specifically would like to thank Dr. Jeffrey Pigott for his assistance with SEM, and Dr. Tae-Kyong John Kim for his assistance with XPS.

## REFERENCES

1. Wang, N. X. & von Recum, H. A. Affinity-Based Drug Delivery. Eng. Polym. Syst. Improv. Drug Deliv. 429–452 (2013). doi:10.1002/9781118747896.ch13

2. Muankaew, C. & Loftsson, T. Cyclodextrin-Based Formulations: A Non-Invasive Platform for Targeted Drug Delivery. Basic Clin. Pharmacol. Toxicol. 122, 46–55 (2018).

3. Ortiz Mellet, C., García Fernández, J. M. & Benito, J. M. CHAPTER 5. Cyclodextrins for Pharmaceutical and Biomedical Applications. (2013). doi:10.1039/9781849737821-00094

4. Thatiparti, T. R., Shoffstall, A. J. & von Recum, H. A. Cyclodextrin-based device coatings for affinity-based release of antibiotics. Biomaterials 31, 2335–2347 (2010).

5. Haimhoffer, Á. et al. Cyclodextrins in drug delivery systems and their effects on biological barriers. Sci. Pharm. 87, (2019).

6. Learn, G. D., Lai, E. J. & von Recum, H. A. Cyclodextrin polymer coatings resist protein fouling, mammalian cell adhesion, and bacterial attachment. bioRxiv 21–24 (2020). doi:10.1101/2020.01.16.909564

7. Sobocinski, J. et al.. Mussel inspired coating of a biocompatible cyclodextrin based polymer onto CoCr vascular stents. ACS Appl. Mater. Interfaces 6, 3575–3586 (2014).

8. Kersani, D. et al.. Stent coating by electrospinning with chitosan/poly-cyclodextrin based nanofibers loaded with simvastatin for restenosis prevention. Eur. J. Pharm. Biopharm. 150, 156–167 (2020).

9. Torii, S. et al.. Drug-eluting coronary stents: insights from preclinical and pathology studies. Nat. Rev. Cardiol. (2019). doi:10.1038/s41569-019-0234-x

10. Venuti, V. et al.. Combining Raman and infrared spectroscopy as a powerful tool for the structural elucidation of cyclodextrin-based polymeric hydrogels. Phys. Chem. Chem. Phys. 17, 10274–10282 (2015).

11. Zhao, F. et al.. EDTA-Cross-Linked β-Cyclodextrin: An Environmentally Friendly Bifunctional Adsorbent for Simultaneous Adsorption of Metals and Cationic Dyes. Environ. Sci. Technol. 49, 10570–10580 (2015).

12. Fu, A. S., Thatiparti, T. R., Saidel, G. M. & Von Recum, H. A. Experimental studies and modeling of drug release from a tunable affinity-based drug delivery platform. Ann. Biomed. Eng. 39, 2466–2475 (2011).

13. Rohner, N. A., Schomisch, S. J., Marks, J. M. & Von Recum, H. A. Cyclodextrin polymer preserves sirolimus activity and local persistence for antifibrotic delivery over the time course of wound healing. Mol. Pharm. (2019). doi:10.1021/acs.molpharmaceut.9b00144

14. McIlvaine, T. C. A Buffer Solution for Colorimetric Comparison. J. Biol. Chem. 49, 183–186 (1921).

15. Carbonate-Bicarbonate Buffer (pH 9.2 to 10.6). AAT Bioquest, INC (2019). Available at: https://www.aatbio.com/resources/buffer-preparations-and-recipes/carbonate-bicarbonate-buffer-ph-9-2-to-10-6. (Accessed: 21st August 2020)

16. Woodard, L. N. & Grunlan, M. A. Hydrolytic Degradation and Erosion of Polyester Biomaterials. ACS Macro Lett. 7, 976–982 (2018).

17. Caldera, F., Tannous, M., Cavalli, R., Zanetti, M. & Trotta, F. Evolution of Cyclodextrin Nanosponges. Int. J. Pharm. 531, 470–479 (2017).

18. Fu, A. S. Affinity-based Delivery and Reloading of Doxorubicin for Treatment of Glioblastoma Multiforme. (Case Western Reserve University, 2013).

